# Computational correction of cross-contamination due to exclusion amplification barcode spreading

**DOI:** 10.1101/176537

**Authors:** Anton JM Larsson, Geoff Stanley, Rahul Sinha, Irving L. Weissman, Rickard Sandberg

## Abstract

Recent advances in sequencing technology have considerably increased the throughput and decreased the cost of short-read sequencing by using high-density patterned flow cells. However, high rates of cross-contamination between multiplexed libraries have been observed on data from these machines, likely due to index-switching during exclusion amplification^1,2^. Here, we demonstrate that a computational correction procedure based on the Sylvester equation removed 80-90% of the false positive expression signal and eliminated spurious clustering. The computational correction procedure can therefore be used to rescue aspects of affected sequence data so that researchers can take advantage of cost-effective sequencing on patterned flow cells.

## Introduction

The massive parallel sequencing capabilities of contemporary sequencers have opened up for large-scale multiplexing of sequencing libraries. In the field of single-cell sequencing^3^, it is common to simultaneously sequence hundreds or thousands of sequencing libraries in the same lane of Illumina sequencers^4-6^, enabling unbiased single-cell analyses of complex tissues^7^ or combinatorial perturbation experiments^8,9^. The integrity of each libraries has thought to be mostly affected by experimental consideration before the construction of complete sequencing libraries, such as avoiding doublets or cross-well contamination. New methods are often evaluated by cross-species experiments, where e.g. human and mouse cells are mixed prior to cell capture and library construction, to enable a quantitative readout of library cross-contamination^5,6^.

Cross-library contamination can also occur during the sequencing itself^2^, and newer Illumina sequencers (e.g. HiSeq 3000, 4000 and NovaSeq) that utilize exclusion amplification (ExAmp) seem particular vulnerable to index-switching caused by free index primers^1^, causing up to 5 or 10% of cross-contaminated reads under particular library construction procedures^1^. These levels of cross-contamination caused several spurious biological interpretations including the artefactual separation of single-cell transcriptomes into sub-types. As free index primers are the source of the cross-contaminating signal, experimental care in removing adapters prior to sequencing becomes important. However, even lower cross-contamination levels however might be problematic for certain applications such as lineage reconstruction from single-cell DNA sequencing, single-cell DNA mutations^10^, tumour mutations^11^ and single-cell allelic expression^12^. Libraries where a single swapping event can lead to the misassignment of a transcript are most affected and include single-cell libraries multiplexed by combinatorial barcoding with i5 and i7 Nextera primers

In this manuscript, we introduce a computational correction procedure inspired by the index-switching observed in hematopoietic stem cell data multiplexed by combinatorial Nextera barcoding^1^. We demonstrate that the correction removed the vast majority of the cross-contamination signal, and that corrected expression data no longer cluster by index, and therefore rescue already sequenced libraries with high levels of index-swapping.

## Results

Since index switching rarely affects both ends of a fragment^1^, the spreading retains information about its source. In each well, the spread counts can be considered a linear combination of the true signal spreading along the columns (among cells sharing an i5 index) and rows (among cells sharing an i7 index) of the library plate through index switching (**Figure 1A**). When the spreading problem is formulated in this manner a method based on linear algebra can be used to find the true counts in the data, given an assumption regarding the rate of the spread. The observed count matrix *C* for each gene can be expressed as the column spread (*AX*) plus the row spread (*XB*), resulting in the equation *AX*+*XB*=*C*. This type of matrix equation is called a Sylvester equation, which is well studied in the mathematical field of control theory, and can be solved by existing algorithms (Methods).

**Figure 1.**
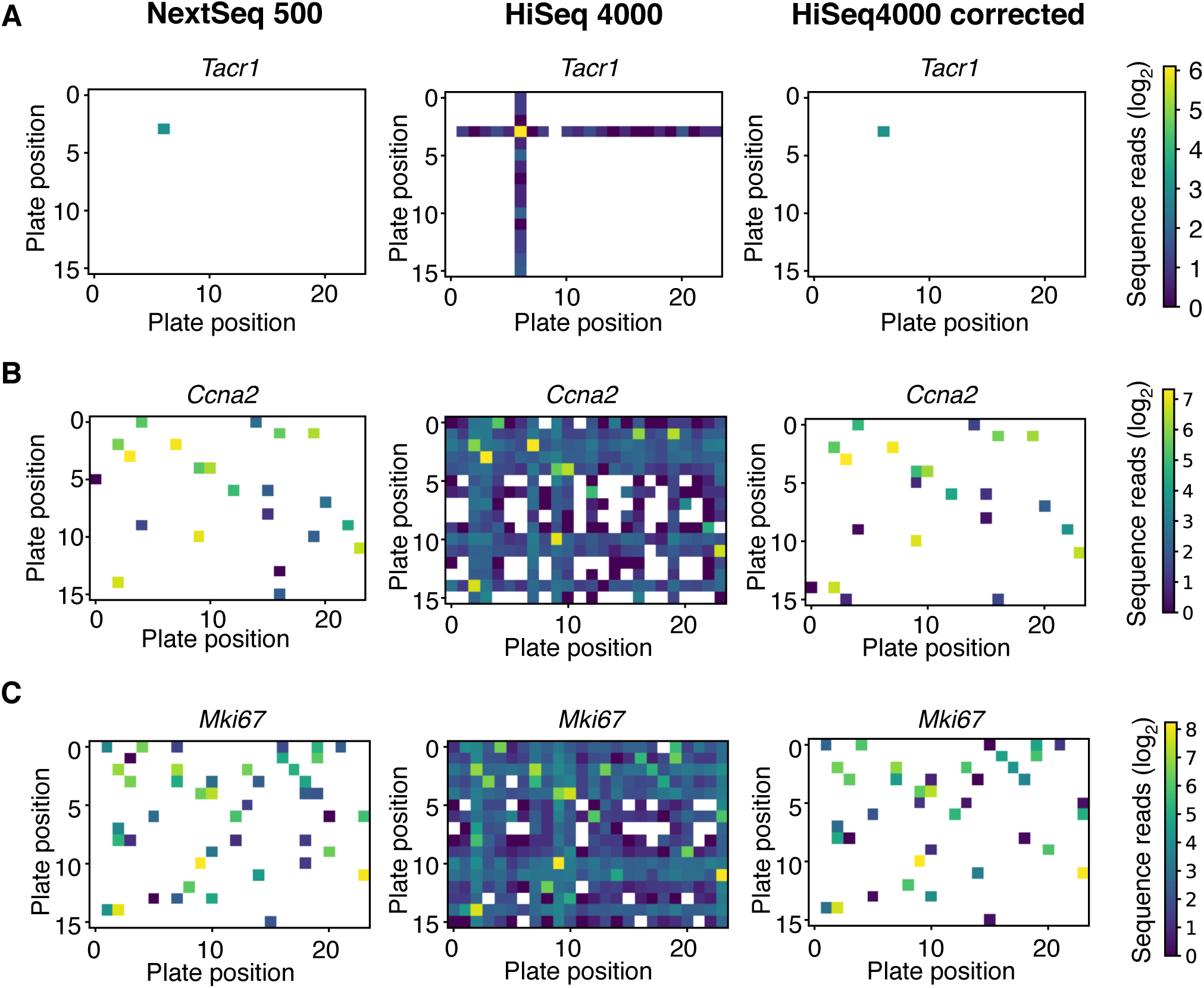
Illustrating index-switching and the computational correction. (**A-C**) A series of heatmaps showing the counts (log scale) of Tacr1 (A), Ccna2 (B) and Mki67 (C). Left: Read counts when sequenced on unaffected NextSeq 500; Middle: Read counts when sequenced on affected HiSeq 4000; Right: Read counts when sequenced on affected HiSeq 4000 after correction.

In order to directly test the correction procedure, we applied it on affected scRNA-seq data from hematopoietic stem cells (HSCs)^1^ which was sequenced on the HiSeq 4000. The correction was then compared to data from the exact same libraries sequenced on the NextSeq 500, which do not have the reported problem of index switching. We assumed that the spread was uniform across primers and assigned each combination of primers an equal rate of spreading (0.01, determined empirically). Then we constructed matrices of the read count for each gene where each entry corresponds to a specific index primer combination, and solved the Sylvester equation. Thereafter we applied a cut-off of 5 reads to remove false positives. Note that simply applying a cut-off to the uncorrected dataset is not sufficient to recover the true expression since the cross-contamination spread signal has a large dynamic range.

We visualized the read counts over the 384-well library plates for specific genes, representative of index-spreading from a single cell source (Tac1 gene in **Figure 1A**) or multiple cells (Ccna2 and Mki67, **Figure 1B-C**). Remarkably, the computational correction recovers the main shape of the true read counts even for genes with true expression in large numbers of cells (**Figure 1B-C**). Comparing the corrected reads counts to the unaffected NextSeq data, we observed occasional loss of expression that was detected on the NextSeq and that the corrected data maintained low expression for cells where none was detected on the NextSeq.

To systematically quantify the degree to which the correction removes the cross-contamination signal, we defined expression detected in the corrected HiSeq data but not the NextSeq data as false positives and likewise expression only detected on the NextSeq as false negatives. Transcriptome-wide, we observed a massive reduction in false positive expression after performing the correction (**Figure 2A**). On a per-gene basis, the correction removed ∼80% of false positive signals (**Figure 2B**). At the same time, we observed a modest increase in false negatives after the correction (**Figure 2C**). These false positive and negative estimates should be conservative, as discrepancies for low expressed genes can also be attributed to the sequencing variability (incomplete sampling).

**Figure 2.**
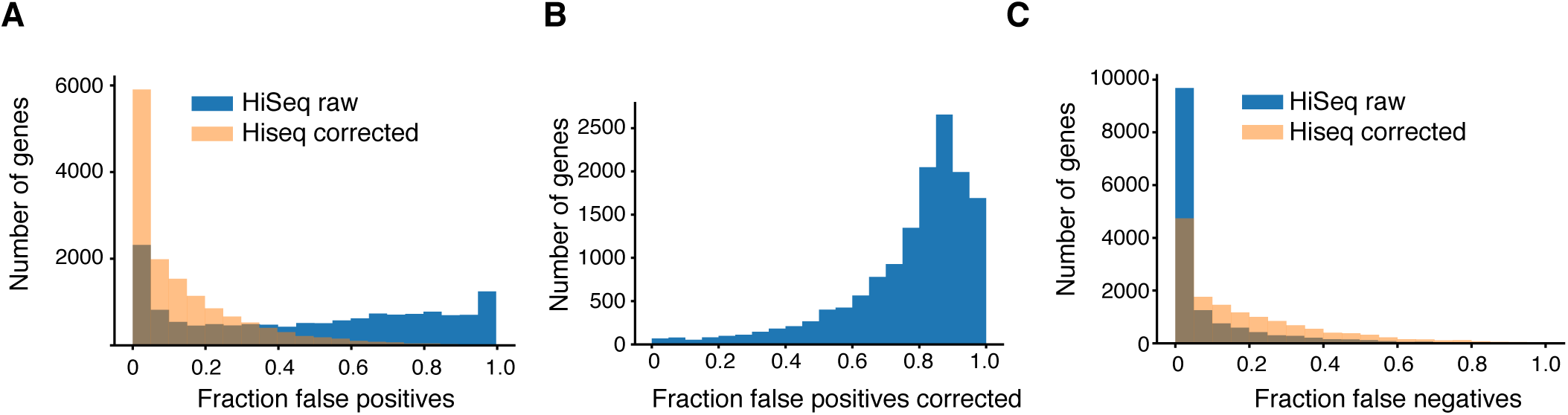
Evaluation of transcriptome-wide correction. (**A**) Histogram of false positive gene expression, defined as gene expression detected on the HiSeq 4000 before (blue) and after correction (yellow) where no expression was detected in the same cell on the NextSeq 500. (**B**) Histogram with fraction false positive expression signals removed per gene. (**C**) Histogram of false negative gene expression defined as no detectable gene expression on the HiSeq 4000 before (blue) and after correction (yellow) where expression was detected in the same cell on the NextSeq 500.

Using robust principal component analysis (rPCA) to cluster individual HSCs by their HiSeq4000 expression levels revealed sub-clusters that corresponded to column and row positions of the cells in the library plates (**Figure 3A-C**). Importantly, clustering by well columns and rows were eliminated after the computational correction of the HiSeq4000 data (**Figure 3D-F**). Thus, the index-spreading observed among the HSCs was sufficient to influence the unbiased clustering of cells. It was also noted that batch effects appeared between different plates of HSCs that were sequenced on the HiSeq4000 data (**Figure 4A**). These batch-effects were also eliminated after the computational correction of index-spreading (**Figure 4B**). We conclude that index spreading can lead to spurious biological interpretations and that the computational correction procedure we introduce was able to remove these artefactual signals in the data.

**Figure 3.**
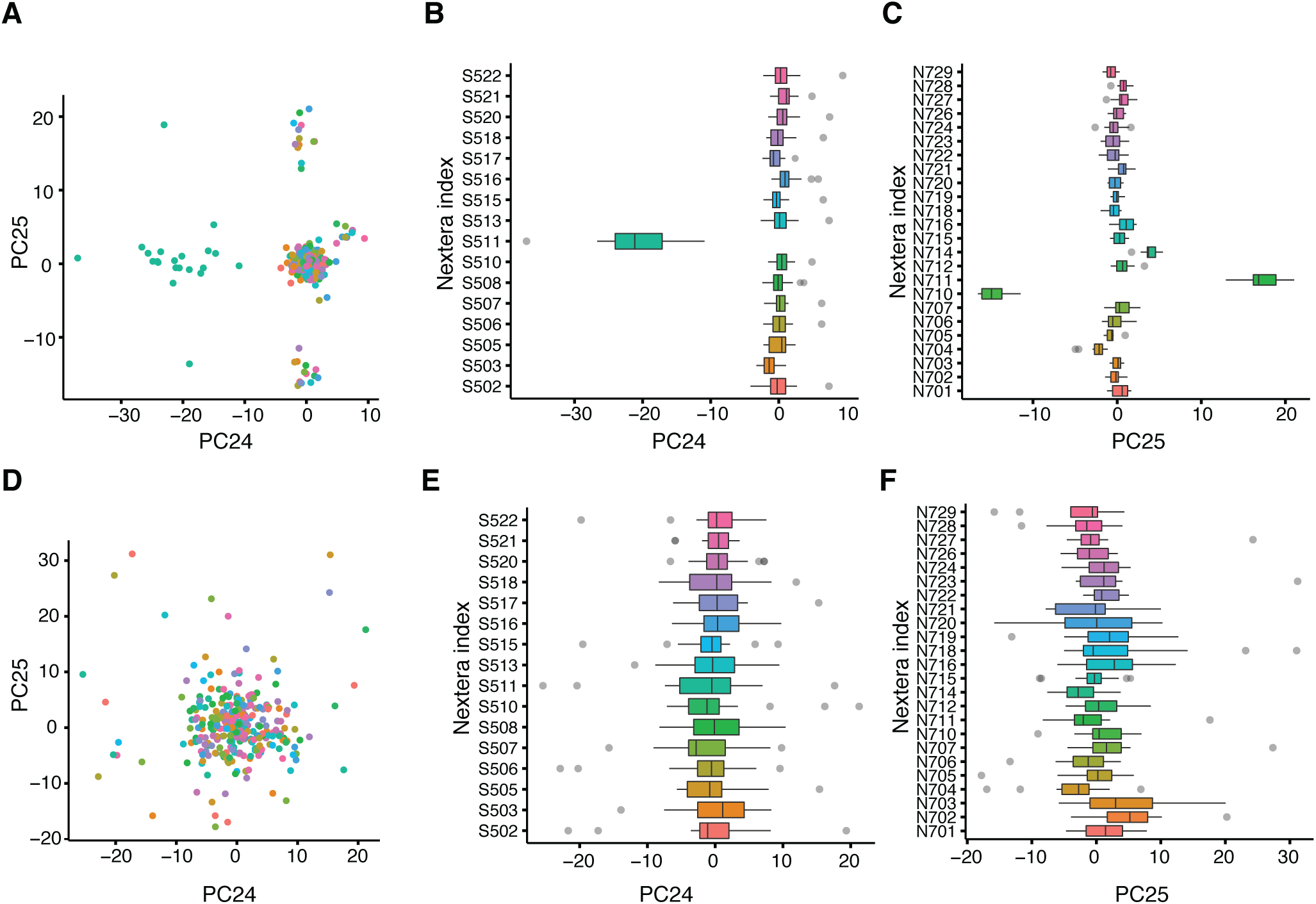
Corrected data rescued spurious clustering. (**A**) Robust PCA (rPCA) analyses of HSCs based on HiSeq4000 sequence counts, coloured by the Nextera i5 index. The rPCA algorithm assign the most prominent outlier observations (i.e. cells) into the last principal components, which corresponded to wells that had been amplified using the same i7 and i5 indices. (**B**) Loadings of cells on PC24, stratified by i5 Nextera primers. (**C**) Loading of cells on P25, stratified by i7 Nextera primers. (**D**) rPCA clustering of HSCs (as in A) for corrected HiSeq4000 sequence counts, colored by Nextera i5 index primer. (**E-F**) As in (C-D) for corrected HiSeq4000 sequence counts.

**Figure 4.**
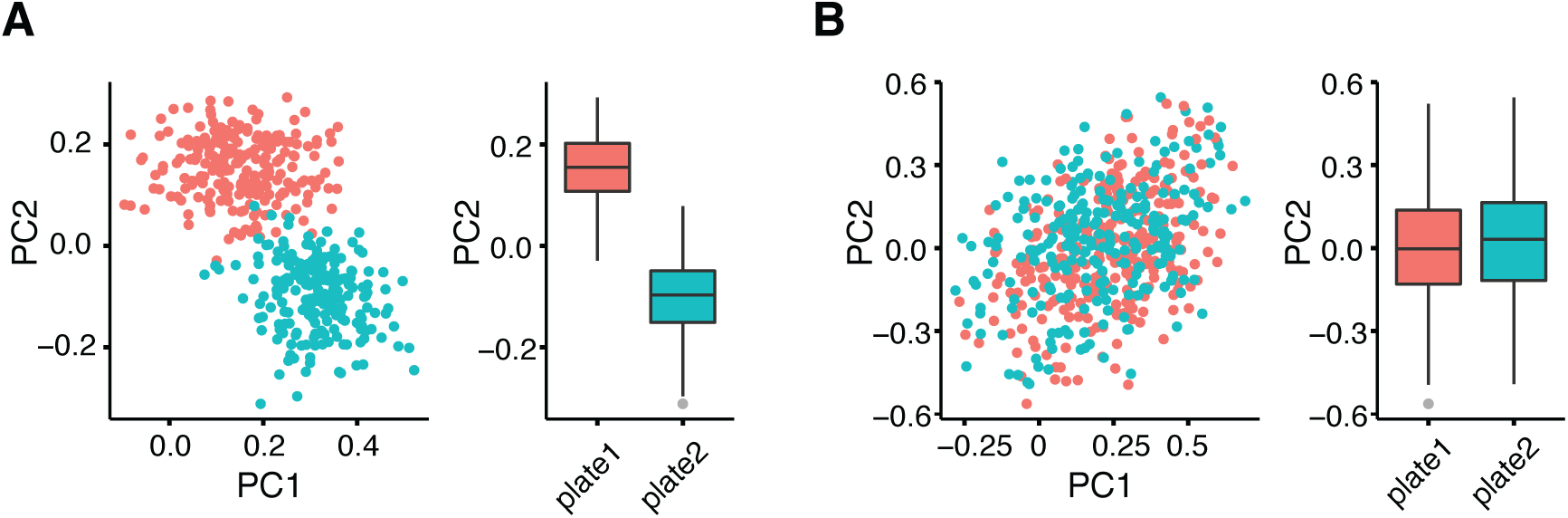
Corrected data removed plate-based clustering. (**A**) PCA analysis of HSCs from two plates of libraries sequenced on HiSeq4000, colored by library plate. Right: PC2 scores separate cells by plates. (**B**) As in (A) for corrected transcriptome data.

## Discussion

We would like to mention that index-switching may be reduced to a large degree during the experimental steps before sequencing through extensive washing. However, it is not clear at this point what steps will need to be taken, how expensive or time-consuming they will be, and to what degree they will reduce the index switching. Furthermore, they may be incompatible with particular experimental protocols, or researchers may want to correct libraries that have already been sequenced. Therefore, the computational correction procedure we introduce will be a critical tool for researchers who take advantage of the throughput and cost-reduction offered by the latest-generation patterned flow cell sequencing machines.

Careful analysis of the concerned data is however recommended before applying this method, since the application of this correction to a dataset which does not suffer from the spreading of signal problem may negatively affect downstream analysis and conclusions. One simple way to identify whether a dataset is affected by the problem is to format the read counts as seen in **Figure 1** and examine genes only highly expressed in a few cells as in **Figure 1A**. Additionally, the rate of switching can be calculated from the number of reads aligning to index combinations not present in the original library, and we suggest that such “blank” wells be included in future libraries sequenced on machines affected by index switching.

## Methods

### Computational correction procedure

The observed read counts in each well can be considered a linear combination of true signal spreading along columns and rows of the plate. Let *A* be a *n* × *n* matrix of the modelled spread along the column of the plate due to the *n* P5 index primers, and B a *m* × *m* matrix of the modelled spread along the row of the plate due to the *m* P7 index primers. Let *X* be an *n* × *m* matrix representing the true counts in each well. Then our observed count matrix *C* for each gene is the column spread (*AX*) plus the row spread (*XB*). We have the resulting equation *AX*+*XB*=*C*. This type of matrix equation is called a Sylvester equation, which is well studied in the mathematical field of control theory. Algorithms to solve this kind of equation are available in most programming languages. In addition, it is worth noting that for our purposes the matrices *A* and *B* which we use to model our spread are always constructed such that *X* has a unique solution. In this study, we used the scientific python (scipy) implementation to solve the Sylvester equation. The rate of spreading along the rows and columns was empirically determined to 0.01, considering the ratio of counts observed in the contaminated wells to the counts in the source well.

### Single-cell RNA-seq data

In this study, we analysed single-cell RNA-sequencing data generated from mouse hematopoietic stem cells using the Smart-seq2 protocol^13^, with important modifications such as the omission of a PCR step after library pooling (see Sinha, Geoff et al.^1^ for details). Library pools were loaded on either an Illumina HiSeq 4000 or NextSeq 500. The processing of sequenced reads into expression matrices were carried out as detailed^1^.

### Clustering analyses of cells and plates

Cells were analyzed using log-transformed counts per million. To demonstrate index clustering, 308 cells from the first library plate (mHSC-plate1), excluding cycling and low-quality cells, were analyzed with rPCA with 25 PCs called. Plot axes labeled “PC*i*” were cells annotated with default PC scores; plot axes labeled “PC*i*.score” were cells scored by sums of the top 30 genes positively- and negatively-correlated to the *i*^th^ principal component. The implementation of Robust PCA used places heterogeneity associated with small subgroups of cells in the final 2-3 PCs^14,15^. To demonstrate separation of sequencing plates, 430 non-cycling cells that passed quality threshold from two HiSeq 4000 sequencing libraries (mHSC-plate1 and mHSC-plate2) were analyzed with rPCA called with 25 PCs.

### Code availability

A Jupyter notebook containing the complete workflow used in this manuscript, including the recreation of figures, will be available at Github (https://github.com/sandberg-lab/spreading-correction) and is also included as an html page in supplement.

## ACKNOWLEDGEMENTS

This work was supported by grants from the Swedish Research Council and the Vallee Foundation.

## AUTHOR CONTRIBUTIONS

A.L. conceived the idea, developed the method, performed the analyses and wrote the manuscript. G.S. performed analyses, R.S. contributed single-cell RNA-seq data, R.S. supervised the project and wrote the manuscript.

## COMPETING FINANCIAL INTERESTS

The authors declare no competing financial interests.

## References

1. Sinha, R. et al. Index Switching Causes ’Spreading-Of-Signal’ Among Multiplexed Samples In Illumina HiSeq 4000 DNA Sequencing. bioRxiv 125724 (2017). doi:10.1101/125724

2. Illumina Inc. Effects of Index Misassignment on Multiplexing and Downstream Analysis.

3. Sandberg, R. Entering the era of single-cell transcriptomics in biology and medicine. Nat. Methods 11, 22–24 (2014).

4. Segerstolpe, Å. et al. Single-Cell Transcriptome Profiling of Human Pancreatic Islets in Health and Type 2 Diabetes. Cell Metab. 24, 593–607 (2016).

5. Macosko, E. Z. et al. Highly Parallel Genome-wide Expression Profiling of Individual Cells Using Nanoliter Droplets. Cell 161, 1202–1214 (2015).

6. Klein, A. M. et al. Droplet barcoding for single-cell transcriptomics applied to embryonic stem cells. Cell 161, 1187–1201 (2015).

7. Zeisel, A. et al. Brain structure. Cell types in the mouse cortex and hippocampus revealed by single-cell RNA-seq. Science 347, 1138–1142 (2015).

8. Adamson, B. et al. A Multiplexed Single-Cell CRISPR Screening Platform Enables Systematic Dissection of the Unfolded Protein Response. Cell 167, 1867–1882.e21 (2016).

9. Jaitin, D. A. et al. Dissecting Immune Circuits by Linking CRISPR-Pooled Screens with Single-Cell RNA-Seq. Cell 167, 1883–1896.e15 (2016).

10. Woodworth, M. B., Girskis, K. M. & Walsh, C. A. Building a lineage from single cells: genetic techniques for cell lineage tracking. Nat. Rev. Genet. 18, 230–244 (2017).

11. Navin, N. et al. Tumour evolution inferred by single-cell sequencing. Nature 472, 90–94 (2011).

12. Reinius, B. et al. Analysis of allelic expression patterns in clonal somatic cells by single-cell RNA-seq. Nat. Genet. 48, 1430–1435 (2016).

13. Picelli, S. et al. Smart-seq2 for sensitive full-length transcriptome profiling in single cells. Nat. Methods 10, 1096–1098 (2013).

14. Gokce, O. et al. Cellular Taxonomy of the Mouse Striatum as Revealed by Single-Cell RNASeq. Cell Rep 16, 1126–1137 (2016).

15. Todorov, V. & Filzmoser, P. An Object-Oriented Framework for Robust Multivariate Analysis. Journal of Statistical Software 32, 1–47 (2009).

